# Linkage mapping reveals loci that underlie differences in *C. elegans* growth

**DOI:** 10.1101/2022.04.25.489412

**Authors:** Joy Nyaanga, Erik C. Andersen

**Affiliations:** Department of Molecular Biosciences, Northwestern University, Evanston, IL 60208, USA; Interdisciplinary Biological Sciences Program, Northwestern University, Evanston, IL 60208, USA

**Keywords:** Developmental growth, *C. elegans*, linkage mapping, QTL

## Abstract

Growth rate and body size are complex traits that contribute to the fitness of organisms. The identification of loci that underlie differences in these traits provides insights into the genetic contributions to development. Leveraging *Caenorhabditis elegans* as a tractable metazoan model for quantitative genetics, we can identify genomic regions that underlie differences in growth. We measured post-embryonic growth of the laboratory-adapted wild-type strain (N2) and a wild strain from Hawaii (CB4856), and found differences in body size. Using linkage mapping, we identified three distinct quantitative trait loci (QTL) on chromosomes IV, V, and X that are associated with variation in body size. We further examined these size-associated QTL using chromosome substitution strains and near-isogenic lines, and validated the chromosome X QTL. Additionally, we generated a list of candidate genes for the chromosome X QTL. These genes could potentially contribute to differences in animal growth and should be evaluated in subsequent studies. Our work reveals the genetic architecture underlying animal growth variation and highlights the genetic complexity of body size in *C. elegans* natural populations.

## Introduction

Precise regulation of final body size is essential to the development and fitness of organisms. Although a larger body size can increase competitive advantages, it also requires added time and nutrients to develop (Hone and Benton 2005). For this reason, mechanisms that control developmental growth rate and ultimate body size are likely under strong natural selection.

The robustness and precision with which animal development is choreographed is still not well understood. Developing systems coordinate the organization and interaction among cells, tissues, and organs at high reproducibility even in the presence of genetic and environmental perturbations. The early developmental biologist C.H. Waddington coined the term “canalization” to describe this biological robustness (Waddington 1942). Developmental canalization has been widely studied in human growth (Tanner 1963; Cameron and Bogin 2012; Desmond and Casale 2017). To achieve an adult height within the normal range, shorter individuals tend to undergo accelerated growth whereas taller individuals experience a decreased rate of growth (Hector and Nakagawa 2012). In this way, the growth curves of individuals, though variable, converge on a narrow range.

To study the phenomenon of organismal size uniformity, considerable precision and throughput is needed, which can be a challenge when working with multicellular organisms. The nematode *Caenorhabditis elegans* is a powerful model organism to study developmental growth because it has a quick generation time, produces large numbers of genetically identical offspring, and is easily cultured in controlled laboratory conditions (Wood 1988). Furthermore, *C. elegans* post-embryonic development is well characterized and marked by four larval-stage transitions (molts) that separate the *C. elegans* life cycle into five distinct stages: four larval stages (L1-L4) and adult (Singh and Sulston 1978). The timing of these molts determines the completion of stage-specific development (Zaidel-Bar *et al*. 2010; Monsalve *et al*. 2011), underscoring the importance of developmental growth regulation in *C. elegans*.

We can leverage *C. elegans* natural genetic diversity to connect phenotypic differences to genetic variants (Evans *et al*. 2021; Andersen and Rockman 2022). Two particular strains of interest are the laboratory-adapted wild-type strain, N2, and a wild strain from Hawaii, CB4856. The genetic diversity between these two strains was shown to underlie multiple phenotypic differences, including aggregation behavior, life history traits, and gene expression (Evans *et al*. 2021). Recombinant inbred lines constructed from crosses between the N2 and CB4856 strains each have unique variants derived from each parental background. Performed at a large scale, these populations of recombinant individuals are a powerful tool to identify genomic regions that are correlated with phenotypic variation. Mapping the natural variation underlying phenotypic differences allows for the dissection of genetic networks involved in important biological processes. Many others have taken this approach to study the genetic underpinnings of complex traits (Evans *et al*. 2021; Andersen and Rockman 2022).

To characterize the genetic basis for variation in body size and growth in *C. elegans*, we first performed a longitudinal study of post-embryonic growth in N2 and CB4856 animals. Although we observed similar patterns in overall growth dynamics, we also noticed small differences in body size at individual time points across development. To study these differences in size, we used linkage mapping to identify three distinct QTL that influence animal size variation. We further assessed each QTL independently using chromosome substitution strains and near-isogenic lines. Doing so, we validated the chromosome X QTL and identified promising candidate genes that could contribute to the differences in size between the N2 and CB4856 strains. Our work provides a framework for future studies to investigate the genetic mechanisms controlling developmental growth and body size in natural populations of *C. elegans*.

## Results

### Growth dynamics of the N2 and CB4856 strains

To precisely evaluate *C. elegans* growth dynamics, we previously developed a high-throughput growth assay that integrates image-based and flow-based devices to quantify the growth of thousands of animals over developmental time (Nyaanga *et al*. 2021). We used this assay to collect body size measurements of N2 and CB4856 animals over the course of larval development from the L1 stage through the L4 stage. Briefly, populations of 100,000 animals were cultured in flasks in triplicate for each strain. Every hour after feeding, we sampled approximately 300 animals from each flask, collected images, and measured length (TOF) and width (norm.EXT) of sampled animals using the COPAS BIOSORT platform (**S1 Fig** and **S1 File**). From these raw body size measurements, we removed non-animal objects using model-based clustering and generated summary statistics to study population changes (**S2-S3 Figs, S2 File**, see **Methods**). Here, we report mean length and mean width of animals over 51 consecutive developmental time points (**Figure 1**). Overall, we observed little divergence in growth behavior between the two strains. As previously reported, we detected continuous growth punctuated by periods of discontinuous growth rate, resulting in visible shifts in length and width over time. Although growth behavior is consistent in both N2 and CB4856 animals, we observed significant differences in animal length and width at individual time points, particularly early in development (**S4 Fig**). As animals age, we identify fewer time points with significant differences likely because of increased variability in body size across the population. We find that in all instances where we observed a significant differences in animal length, animals from the N2 strain were consistently longer than animals from the CB4856 strain. However, this result was not observed in animal width as we observed time points where the CB4856 strain was wider than the N2 strain and other timepoints with the opposite result.

**Figure 1.**
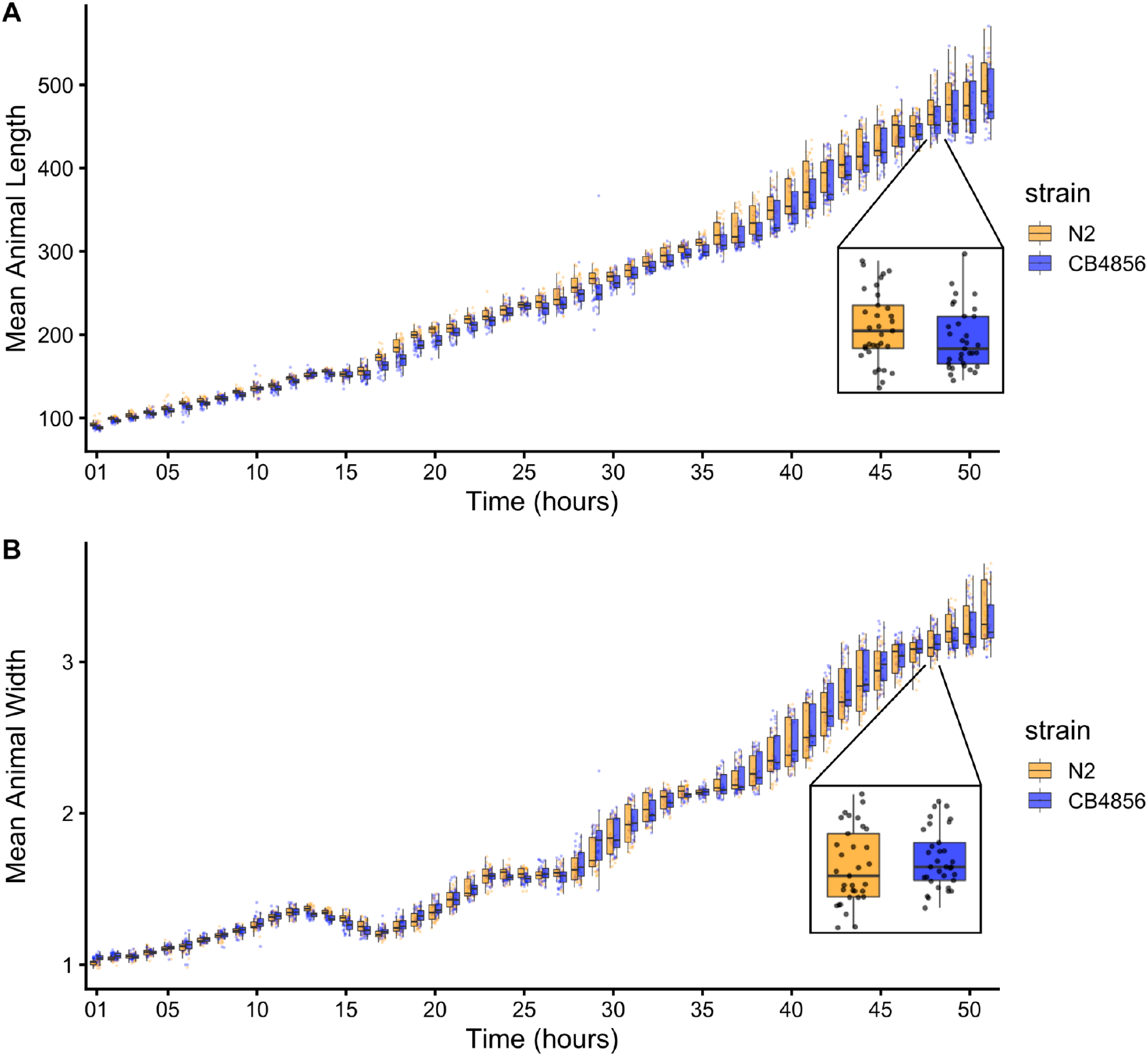
Quantitative measurements of growth for N2 and CB4856 animals. Tukey boxplots of mean length **(A)** and mean width **(B)** for the N2 (orange) and CB4856 (blue) strains over developmental time. The horizontal line in the middle of the box is the median, and the box denotes the 25th to 75th quantiles of the data. The vertical line represents the 1.5 interquartile range. Inset plots magnify mean animal size measurements from hour 48. Each point corresponds to the mean length or mean width of a population of animals in each well.

### Identification of QTL underlying variation in growth

As a complex trait, developmental growth is likely influenced by many genes as well as the interactions among them. To investigate the genetic basis of differences in growth, we assessed the development of a panel of 310 RIAILs derived from a cross between the N2 and CB4856 strains (set 2 RIAILs, see **Methods**). In lieu of collecting measurements throughout development, we used body size as a convenient proxy for developmental progression, where fast growth corresponds to large size and slow growth corresponds to small size. After 48 hours post-L1 arrest, we collected measurements of length and width using a high-throughput fitness assay, and removed wells containing more than 100 or fewer than three animals from downstream processing (**S3 File**, see **Methods**). Doing so, we observed a distribution of both mean length (**Figure 2A**) and mean width (**Figure 2D**) among the RIAILs, indicating that growth rate varies in the strain population. Next, we mapped body length and width separately and obtained three significant QTL (**Figure 2B,E** and **S4 File**). The length-associated QTL on the center of chromosome IV and the width-associated QTL on the center of chromosome V independently explain approximately 5% of the phenotypic variation among the RIAILs. The third QTL on the right arm of chromosome X explains slightly more variation at 8.6% (**Table 1**). Notably, not only did we find distinct QTL for length and width, we also observed QTL with opposite effects on body shape. Strains with the N2 allele at the chromosome IV QTL were longer than strains with the CB4856 allele at this QTL (**Figure 2C**). By contrast, strains with the CB4856 alleles at the chromosome V and X QTL were wider than strains with the N2 allele at these QTL (**Figure 2F**). The identification of distinct QTL for length *vs*. width indicates that body shape is influenced by multiple genetic mechanisms. Additionally, we scanned the genome for interactions between pairs of genomic markers that could affect the phenotypic distribution of length or width in the RIAILs and identified no significant interactions (**S5-S6 Figs**). These data suggest that the three identified loci contain variants that uniquely influence growth rate along multiple axes, where each locus independently affects the longitudinal or circumferential growth of animals.

**Table 1.** Body size QTL.

| Trait | Chromosome | Interval (bp) | Peak | LOD | Variance explained (%) | Effect size |
| --- | --- | --- | --- | --- | --- | --- |
| Length | IV | 6,211,685 - 12,868,784 | 9,392,639 | 3.22 | 5.28 | -0.229 |
| Width | V | 5,371,124 - 12,112,105 | 11,806,498 | 4.54 | 5.59 | 0.236 |
| Width | X | 12,565,734 - 13,173,080 | 12,750,794 | 5.25 | 8.63 | 0.293 |

**Figure 2.**
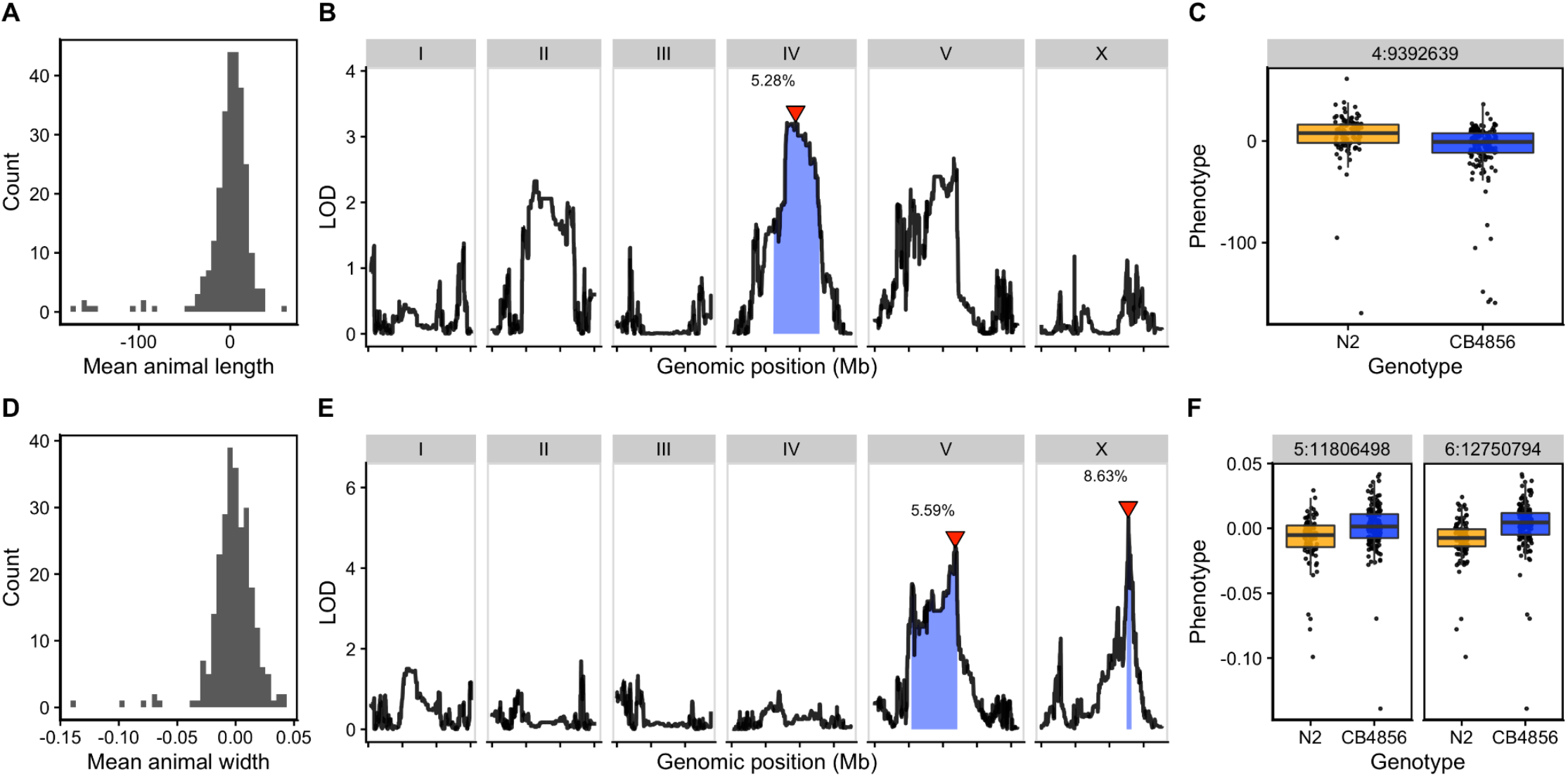
Linkage mapping identifies three QTL associated with body size. Histogram of normalized mean body length **(A)** and mean body width **(D)** of the RIAIL population. **(B, E)** Linkage mapping results for mean body length or mean body width are shown with genomic position (x-axis) plotted against the logarithm of the odds (LOD) score (y-axis). X-axis tick marks denote every 5 Mb. Significant QTL are denoted by a red triangle at the peak marker, and blue shading shows the 95% confidence interval around the peak marker. The percentage of the total phenotypic variance in the RIAIL population that is explained by each QTL is shown above the peak marker. **(C, F)** Tukey box plots show the normalized mean length or width (y-axis) of RIAILs split by genotype at the marker with the maximum LOD score (x-axis). Populations of recombinant strains were grown in independent wells. Each point corresponds to the mean value calculated from each assayed well. Boxes for data from strains with the N2 allele are colored orange, and boxes for data from strains with the CB4856 allele are shown in blue.

### Validation of loci associated with differences in animal size

To validate whether genetic variation between the N2 and CB4856 strains contributes to differences in animal size, we generated chromosome substitution strains (CSSs) for chromosomes IV and V in which the entire chromosome from the N2 strain was introgressed into the CB4856 genetic background and vice versa. We also constructed reciprocal near-isogenic lines (NILs) for the chromosome IV, V, and X QTL (**S5-S6 Files**). These NILs contain a small genomic segment derived from one parent strain introgressed into the genetic background of the other parent strain. We then measured the length and width of animals after 48 hours of growth and calculated statistical significance in a pairwise manner for each strain (**S7-S8 Files**, see **Methods**). For the chromosome IV and V QTL, we were unable to recapitulate the results observed in the linkage mapping (**S7**-**S8 Fig** and **S9 File**). These two QTL each explain only 5% of the total phenotypic variation among the RIAILs and have the smallest effect sizes among the three detected QTL (**Table 1**). The inability to validate these QTL suggests a complex genetic architecture that cannot be explained by isolating these loci using CSSs and NILs, or a lack of power to detect differences driven by these QTL in the CSSs and NILs. By contrast, we successfully validated the chromosome X QTL by observing that genotype significantly contributed to differences in body width of NILs (**Figure 3** and **S10 File**). The strain with the CB4856 allele on chromosome X crossed into the N2 genetic background was significantly wider than the N2 strain (Tukey’s HSD, *p-*value = 1.29*e*^-10^). Similarly, the strain with the N2 chromosome X region introgressed into the CB4856 genetic background was significantly thinner than the CB4856 strain (Tukey’s HSD, *p*-value = 1.29*e*^-10^). These results confirmed that genetic variation between the N2 and CB4856 strains on chromosome X contributes to the difference in body width between these strains.

**Figure 3.**
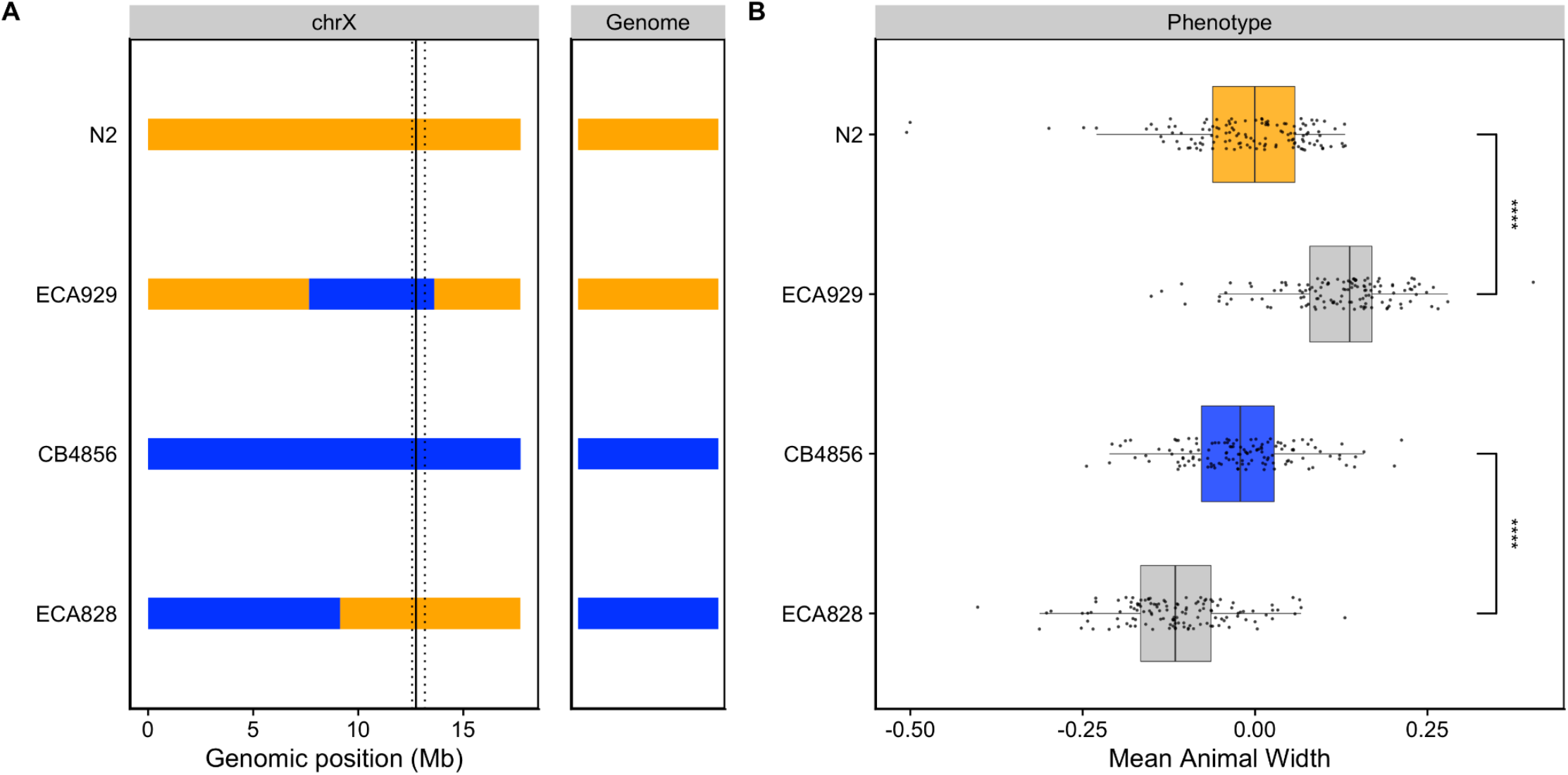
NILs validated the chromosome X QTL. **(A)** Strain genotypes are displayed as colored rectangles (N2: orange, CB4856: blue) for chromosome X (left) and in general for the rest of the chromosomes (right). The solid vertical line represents the peak marker of the QTL. The dashed vertical lines represent the confidence interval. **(B)** Residual mean animal width (x-axis) is plotted as Tukey box plots against strain (y-axis). Each point corresponds to the mean width of a population of animals from each well. The boxes for the parental strains are colored: N2, orange; CB4856, blue. Statistical significance was calculated by Tukey’s HSD (**** = *p-*value < 0.0001).

### Identification of candidate genes in the chromosome X QTL

To identify candidate genes that could underlie variation in body width, we investigated the genes in the chromosome X interval in the N2 strain. We found 151 genes present in this interval and eliminated 96 genes that had no genetic variation in the CB4856 strain (**Table 2, File S11**). Of the remaining 55 genes, 18 have genetic variation in the amino-acid sequence of a protein (protein-coding variation), and 34 have genetic variation that is not protein-coding (non-coding variation). However, protein-coding variation is just one way in which genetic variation can cause phenotypic variation. We also considered instances where genetic variation causes a change in gene expression. Using an expression QTL (eQTL) dataset that mapped expression differences in another panel of RIAILs (set 1) derived from N2 and CB4856 (Rockman and Kruglyak 2009; Evans and Andersen 2020), we identified five genes with eQTL that map to our region of interest. Additionally, we found 17 other genes outside this genomic interval with eQTL that map to this interval, resulting in a total of 72 candidate genes, none of which are located within a hyper-divergent region (Lee *et al*. 2021).

**Table 2.** Genes in QTL interval for chromosome X.

| No variation<sup>a</sup> | Protein-coding variation and/or eQTL<sup>b</sup> | Non-coding variation<sup>c</sup> | Other eQTL that map to interval<sup>d</sup> | Total |
| --- | --- | --- | --- | --- |
| 96 | 21 | 34 | 17 | 168 |

<sup>a</sup>Genes within genomic interval with no genetic variation
<sup>b</sup>Genes within genomic interval with protein-coding variation and/or an eQTL that maps to this interval
<sup>c</sup>Genes within genomic interval with non-coding variation and no eQTL that maps to this interval
<sup>d</sup>Genes outside genomic interval with eQTL that maps to this interval

To further narrow our list of genes, we inspected the functional descriptions and gene ontology (GO) annotations for the 72 candidate genes. When considering the 21 genes with protein-coding variation and/or eQTL, one candidate (*ppk-3)* stood out. *ppk-3*, or phosphatidylinositol phosphate kinase 3, is an ortholog of the mammalian PIKfyve. These kinases play important roles in cell communication and membrane trafficking (Gillooly *et al*. 2001). Notably, mutations in *ppk-3* are responsible for a range of developmental defects, including embryonic lethality, developmental arrest, and larval growth delay (Nicot *et al*. 2006). Investigating the sequence read alignments of the N2 and CB4856 strains at the *ppk-3* locus using the Variant Browser on CeNDR (Cook *et al*. 2017), we observed a missense variant in the second exon predicted to encode a serine-to-threonine substitution (S43T). Although this variant is not in a predicted functional domain, it could alter protein function thereby contributing to the observed phenotypic difference. Aside from *ppk-3*, we identified two additional candidate genes when assessing the functional description for the 34 genes with non-coding genetic variation. The first, *nhr-25*, encodes a nuclear hormone receptor orthologous to Ftz-F1 in *Drosophila* and is required for proper molting and developmental control (Asahina *et al*. 2000). We observed a splice-site variant in the *nhr-25* locus that could disrupt proper RNA splicing. Interestingly, disruption of *nhr-25* often causes embryonic arrest; however, mutants that survive hatching display a squat body stature (Dpy phenotype), suggesting that *nhr-25* could play a role in body size and shape (Gissendanner and Sluder 2000). The second, *bcat-1*, encodes a branched-chain amino acid aminotransferase that was shown to be required for normal embryonic and larval development (Maeda *et al*. 2001). In the *bcat-1* locus, we found a variant in an intron and in the 3’ untranslated region. Together, these results suggest that one or more genes on chromosome X are candidates that need additional study to explain the variation that we observe in animal growth.

## Discussion

Here, we investigated larval growth of N2 and CB4856 animals from the L1 stage to the L4 stage. Although we observed similarities in the dynamics of growth, we also saw differences in the size of animals across developmental time. We used linkage mapping to investigate these differences and identified three small-effect QTL associated with variation in body size. Two QTL underlie variation in animal width, and a single non-overlapping QTL contributes to differences in animal length. Using NILs, we validated the width-associated QTL on chromosome X and identified candidate genes that could underlie variation in width. Taken together, our results demonstrate the power of leveraging natural genetic variation to examine the genetic architecture of complex traits such as body size and shape.

### A complex genetic architecture underlies differences in body size

As a complex life history trait, developmental growth could be influenced by several loci (Houle 1992). In this study, we report three size-associated QTL. Strikingly, we find loci that decouple components of body size, revealing a complex genetic system that influences growth along different axes of the body. Evidence for genetically separate modules underlying distinct aspects of a single trait has been observed in studies of *C. elegans* behavioral patterns where linkage mapping studies using a panel of RIAILs (set 1, (Rockman and Kruglyak 2009)) identified distinct loci underlying separate aspects of a response to thermal stimuli (Ghosh *et al*. 2015). Here, we identify distinct QTL for length and width, suggesting that different genetic mechanisms control animal growth along the length *vs*. width directions. This finding is particularly interesting given the differences in general growth dynamics of length compared to width that we found here (**Figure 1**) and previously (Nyaanga *et al*. 2021) a simultaneous increase in length and decrease in width at the transition between larval stages.

The results of the linkage mapping experiment identified two broad peaks on chromosomes IV and V associated with length and width respectively, as well as a narrow peak on chromosome X for width. Although we successfully validated the width-associated QTL on chromosome X (**Figure 3**), we were unable to validate the other two QTL (**S6** and **S7 Figs**). Our inability to recapitulate the results observed in the linkage mapping might be driven by several factors. First, many loci spread across the genome could underlie variation in body size. Under this polygenic model, any region can harbor variants driving our observed phenotypic difference through additive and/or non-additive effects. The contribution of polygenicity to phenotypic variance has previously been explored in *C. elegans*. Studies of fertility and body size in the *C. elegans* multiparental experimental evolution (CeMEE) panel found that a significant fraction of phenotypic variance, nearly 40% for fertility, can be explained by polygenicity (Noble *et al*. 2017). Second, the intervals could contain QTL of opposing effects, making it difficult to recapitulate the results observed in the mapping using NILs. Notably, researchers have observed patterns of polygeny and antagonistic-effect loci when investigating *C. elegans* growth and reproduction in nickel stress (Bernstein *et al*. 2019). Third, it is possible that the QTL effects are smaller than 5% and we are underpowered to detect differences driven by these QTL in the CSSs and NILs.

### Candidate genes for variation in body size

Genetic variants underlying complex traits are often elusive (Rockman 2012; Evans *et al*. 2021). Ultimately, when searching for QTL, we aim to identify genes contributing to the variation in phenotypes among individuals. Here, we identified candidate genes located in the interval of the chromosome X QTL (**Table 2** and **S11 File**). However, complex traits, such as body size, are likely affected by many genes. For example, recent studies of human genetic variation using data from 5.4 million individuals report finding over 12,000 independent loci associated with height (Yengo *et al*. 2022). In the laboratory strain of *C. elegans*, we know many loci that quantitatively affect body size and shape. Mutations in these genes span various classes, including abnormal pharyngeal pumping (Eat), egg-laying defective (Egl), uncoordinated (Unc), abnormal dauer formation (Daf), and several cuticle and body shape classes (Dpy, Lon, Sma, Rol, Sqt) (Mörck and Pilon 2006; So *et al*. 2011; Cho *et al*. 2021). The polygenic nature of complex traits is a recognized barrier in identifying the genes contributing to phenotypic variation in a population (Boyle *et al*. 2017; Bernstein *et al*. 2019). However, we believe that molecular analysis of loci that underlie variation in development-associated traits in natural populations of *C. elegans* is essential to deciphering the evolutionary significance of developmental canalization.

### Comparison with previous QTL studies of *C. elegans* growth

Our mapping results both recapitulate and expand upon previous QTL studies of growth in *C. elegans*. Previously, median body length of mixed-stage animals was mapped using the same panel of RIAILs (set 2) (Andersen *et al*. 2015). A single small-effect (5.7%) QTL in the center of chromosome IV was found, consistent with our findings. Also in this study, the authors mapped median body width (norm.EXT) to three QTL on chromosomes III, IV, and X. We detected an overlapping genomic region on chromosome X in our current study. The failure to recapitulate the other QTL is likely caused by differences in the body size assay as the previous study measured mixed-stage animals and we focused on synchronized L4 animals. Additionally, others have mapped variation in animal length for a collection of N2xCB4856 introgression lines at 48 hours after L1 arrest (Snoek *et al*. 2014). Here, investigators found five separate QTL on chromosome IV affecting body size. This result suggests the presence of several independent loci on chromosome IV each contributing to variation in length. Further investigation is necessary to determine whether the overlapping genomic region detected in our current study is in fact a separate locus that independently contributes to variation in animal length. Most recently, a group using a N2xCB4856 RIL population identified 18 QTL influencing various body-size traits at a range of temperatures, with the majority clustering on chromosome X (Maulana *et al*. 2022). This work not only demonstrates the genetic complexity underlying body-size phenotypes but also suggests the presence of co-regulatory loci underlying plasticity. *C. elegans* gives investigators a powerful system to understand the genetic mechanisms that shape growth and environmental sensitivity in natural populations.

## Materials and methods

### Strains

Animals were grown at 20°C on 6 cm plates of modified nematode growth media (NGMA), containing 1% agar and 0.7% agarose seeded with *E. coli* OP50 bacteria. Recombinant inbred advanced intercross lines (RIAILs) used for linkage mapping were constructed previously (Andersen *et al*. 2015). The construction of chromosome substitution strains (CSSs) and near-isogenic lines (NILs) used for validation is detailed below. Strains are available upon request.

### High-throughput growth assay

Measurements of body size and fluorescence were measured as previously described (Nyaanga *et al*. 2021). Briefly, the N2 and CB4856 strains were propagated for three generations, bleach-synchronized, and titered at a concentration of 1 embryo per µL into six replicate 500 mL flasks for a final volume of 25 mL. The following day, arrested L1s were fed HB101 food at a final concentration of OD20 in a final flask volume of 100 mL K medium and HB101 food. Animals were then grown at 20°C with constant shaking. Flasks were sampled each hour beginning one hour after feeding and continuing for 51 consecutive hours. At each hour, animals were sampled from each flask, treated with sodium azide, imaged with an ImageXpress Nano (Molecular Devices, SanJose, CA) and scored using a large-particle flow cytometer (COPAS BIOSORT, Union Biometrica, Holliston MA). The COPAS BIOSORT platform was used to collect measurements of animal length (TOF) and optical extinction (EXT). Normalized optical extinction (norm.EXT) was previously established as a proxy for animal width. The raw data collected were imported and processed using the *easysorter* R package (Shimko and Andersen 2014). Processing removed non-animal objects such as bacterial clumps, shed cuticles, and next generation larval animals from the time-course data using the *mclust* R package (Scrucca *et al*. 2016).

### High-throughput fitness assay for linkage mapping

For RIAIL phenotyping, we used a high-throughput fitness assay previously described (Andersen *et al*. 2015). In brief, populations of each strain were propagated on NGMA plates for four generations after which gravid adults were bleach-synchronized and embryos from each strain were aliquoted at a concentration of 25-50 embryos/μL into 96-well microtiter plates for a final volume of 50 μL K medium. The next day, arrested L1s were fed HB101 bacterial lysate (Pennsylvania State University Shared Fermentation Facility, State College, PA; (García-González *et al*. 2017)) at a final concentration of 5 mg/mL in K medium and grown to the L4 larval stage for 48 hours at 20°C with constant shaking. Animals were then sorted using a COPAS BIOSORT platform during which time animal length and width were collected. Measurements collected by the COPAS BIOSORT were processed and analyzed using the *easysorter* R package (Shimko and Andersen 2014). Well populations of recombinant strains that contained more than 100 or fewer than three individuals were removed from further processing, resulting in an average of 25 independent replicate wells per strain. Differences among strains tested on different days were controlled using a linear model (*animal_size ∼ experiment_date*). In this way, we address only the differences among strains caused by growth and the day-to-day experimental variance is controlled. These residual values are used for plotting.

### Linkage mapping

310 RIAILs (set 2 RIAILs) were phenotyped using the high-throughput assay described above. Linkage mapping was performed for body size traits using the R package *linkagemapping* (www.github.com/AndersenLab/linkagemapping) as previously described (Brady *et al*. 2019). The genotypic data and residual phenotypic data were merged using the *merge_pheno* function with the argument *set = 2*. Quantitative trait loci (QTL) were detected using the *fsearch* function. This function calculates the logarithm of the odds (LOD) scores for each genetic marker and each trait as *-n(ln(1-R^2)/2ln(10))* where R is the Pearson correlation coefficient between the RIAIL genotypes at the marker and trait values (Bloom *et al*. 2013). A significance threshold based on a 5% genome-wide error rate was calculated by permuting the phenotypic values of each RIAIL 1000 times. QTL were identified as the marker with the highest LOD score above the significance threshold. This marker was then integrated into the model as a cofactor and mapping was repeated iteratively until no further significant QTL were identified. Finally, the *annotate_lods* function was used to calculate the effect size of each QTL. 95% confidence intervals were defined by a 1.5-LOD drop from the peak marker.

### Generation of chromosome substitution strains (CSSs) and near-isogenic lines (NILs)

CSSs were generated from a cross of the N2 and CB4856 strains. These strains were crossed and heterozygous hermaphrodite progeny were mated to each parental genotype for four generations followed by three generations of selfing to ensure homozygosity of the genome. For each cross, PCR amplicons for insertion-deletions (indels) on the left and right sides of chromosomes IV and V were used to confirm progeny genotypes and select non-recombinants within the introgressed region (**S1 Text**). CSSs were whole-genome sequenced to confirm their genotypes.

NILs were generated as previously described (Zdraljevic *et al*. 2017, 2019; Evans *et al*. 2018; Brady *et al*. 2019) by either backcrossing a selected RIAIL or NIL for six generations or *de novo* by crossing the parental strains N2 and CB4856 to create a heterozygous individual that was then backcrossed for six generations. PCR amplicons for indel variants were used to track the genomic interval (**S1 Text**). NILs were whole-genome sequenced to verify introgressions.

### Statistical analysis of CSS and NIL results

Growth dynamics for CSSs were tested using a modified version of the high-throughput fitness assay for linkage mapping. Animals were propagated on NGMA plates for two generations before gravid adults were bleach-synchronized and embryos from each strain were aliquoted at a concentration of 1 embryo/μL into 12-well, flat bottom culture plates. After three days, gravid adults were bleach-synchronized and embryos were titered into 96-well microtiter plates at a concentration of 50 embryos/μL for a final volume of 50 μL K medium. The next day, arrested L1s were fed HB101 live bacterial food at a final concentration of OD20. Animals were grown for 48 hours at 20°C with constant shaking and then scored using the COPAS BIOSORT platform as before. The raw data collected were again imported and processed using the *easysorter* R package (Shimko and Andersen 2014). Processing removed non-animal objects such as bacterial clumps, shed cuticles, and next generation larval animals from the time-course data using the *mclust* R package (Scrucca *et al*. 2016). Complete pairwise strain comparisons were performed using the *TukeyHSD* function (R Core Team and Others 2013) on an ANOVA model with the formula *phenotype ∼ strain*. A *p-*value of *p* < 0.05 was used as a threshold for statistical significance. Recapitulation was defined by the significance and direction of effect the CSS or NIL had compared to the parental strains.

## Data availability

The authors state that all data necessary to confirm the conclusions of this work are within the text, figures, and supporting information files. All files and code for analysis and generation of figures and tables are archived on GitHub (https://github.com/AndersenLab/N2CB-growth-manuscript).

## Author contributions

Joy Nyaanga: Conceptualization, Investigation, Methodology, Formal Analysis, Writing - original draft, Writing - review and editing Erik C. Andersen: Conceptualization, Funding acquisition, Investigation, Supervision, Writing - review and editing.

## Acknowledgements

We thank Sophia Gibson and Gaotian Zhang for their help sampling animals during the 51-hour growth assay.

