## Supplemental Figures for "Linkage mapping reveals loci that underlie differences in *C. elegans* growth"

**S1 File. Raw animal growth data.**
Raw growth data collected from the COPAS BIOSORT and processed using the *easysorter* R package to compile information from each well. (CSV)
**S2 File. Pruned animal growth data.**
Processed data from the COPAS BIOSORT following implementation of the *mclust* R package and removal of clusters containing non-animal objects. (CSV)
**S3 File. RIAIL phenotype data.**
Residual phenotypic values for all 310 RIAILs, and parent strains (N2 and CB4856) for each trait (TOF = length and normEXT = width). (TSV)
**S4 File. Linkage mapping results.**
Linkage mapping results with annotated QTL and confidence intervals. (CSV)
**S5 File. CSS/NIL sequence data.**
VCF from the whole-genome sequencing for all the CSSs/NILs in this study. (DOCX)
**S6 File. CSS/NIL genotype data.**
Genotypes for the CSSs/NILs in this study. (CSV)
**S7 File. CSS and NIL raw phenotype data.**
Raw phenotype data for the CSSs and NILs in this study. (CSV)
**S8 File. CSS and NIL pruned phenotype data.**
Pruned phenotype data for the CSSs and NILs in this study. (CSV)
**S9 File. Statistical significance for mean animal length from CSS/NIL assay.**
Pairwise statistical significance for all strains. (CSV)
**S10 File. Statistical significance for mean animal width from CSS/NIL assay.**
Pairwise statistical significance for all strains. (CSV)
**S11 File. List of genes in the chromosome X QTL.**
List of all genes in the chromosome X interval, their functional descriptions and GP annotations, and whether they have variation in the CB4856. (CSV)
**S1 Text. Reagents used to generate CSSs and NILs.** (PDF)

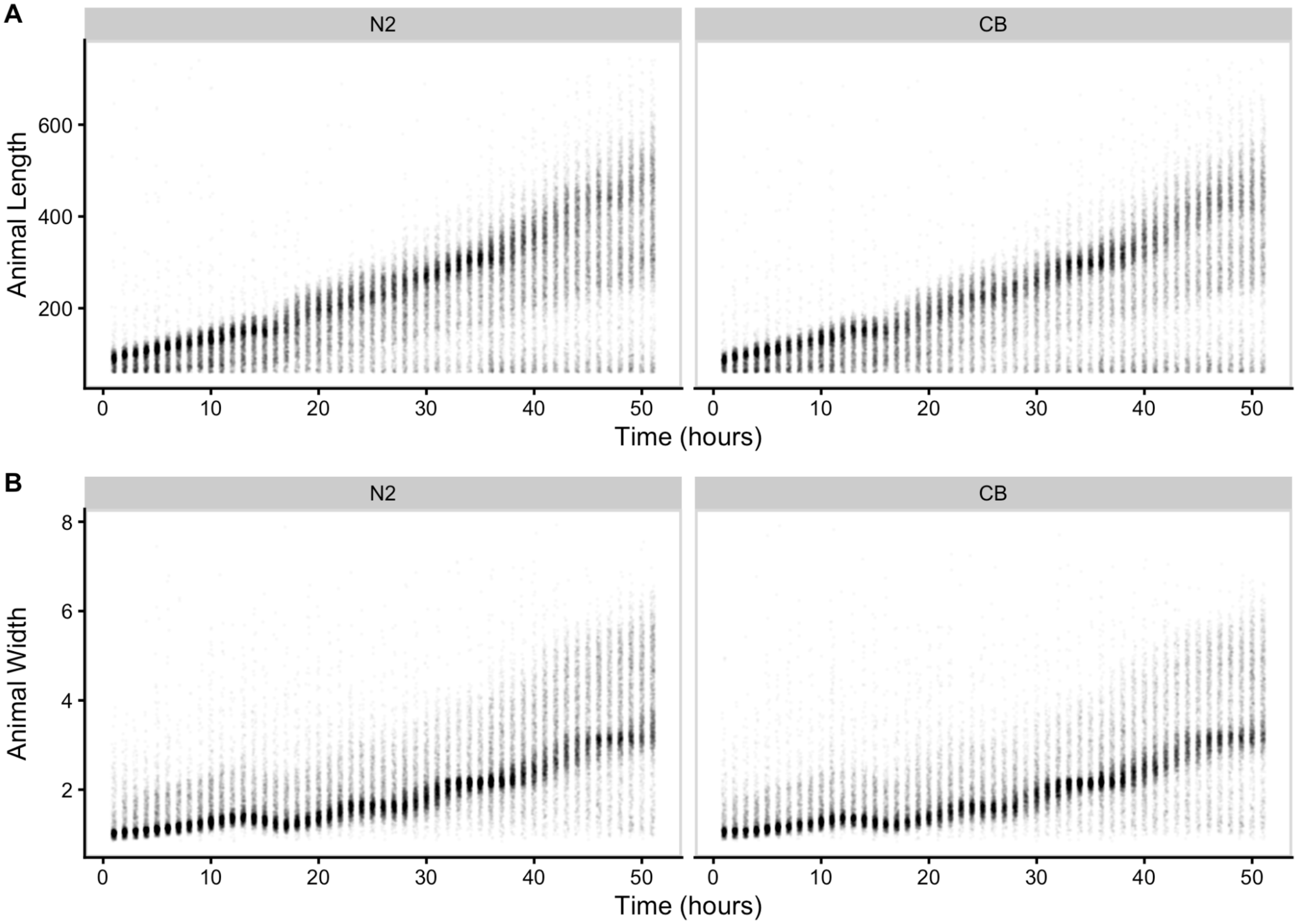


**S1 Fig. Raw measurements of animal size.**
Raw COPAS BIOSORT of animal length (A) and width (B) for N2 and CB4856 objects are shown here. Each point represents an individual object that was measured.

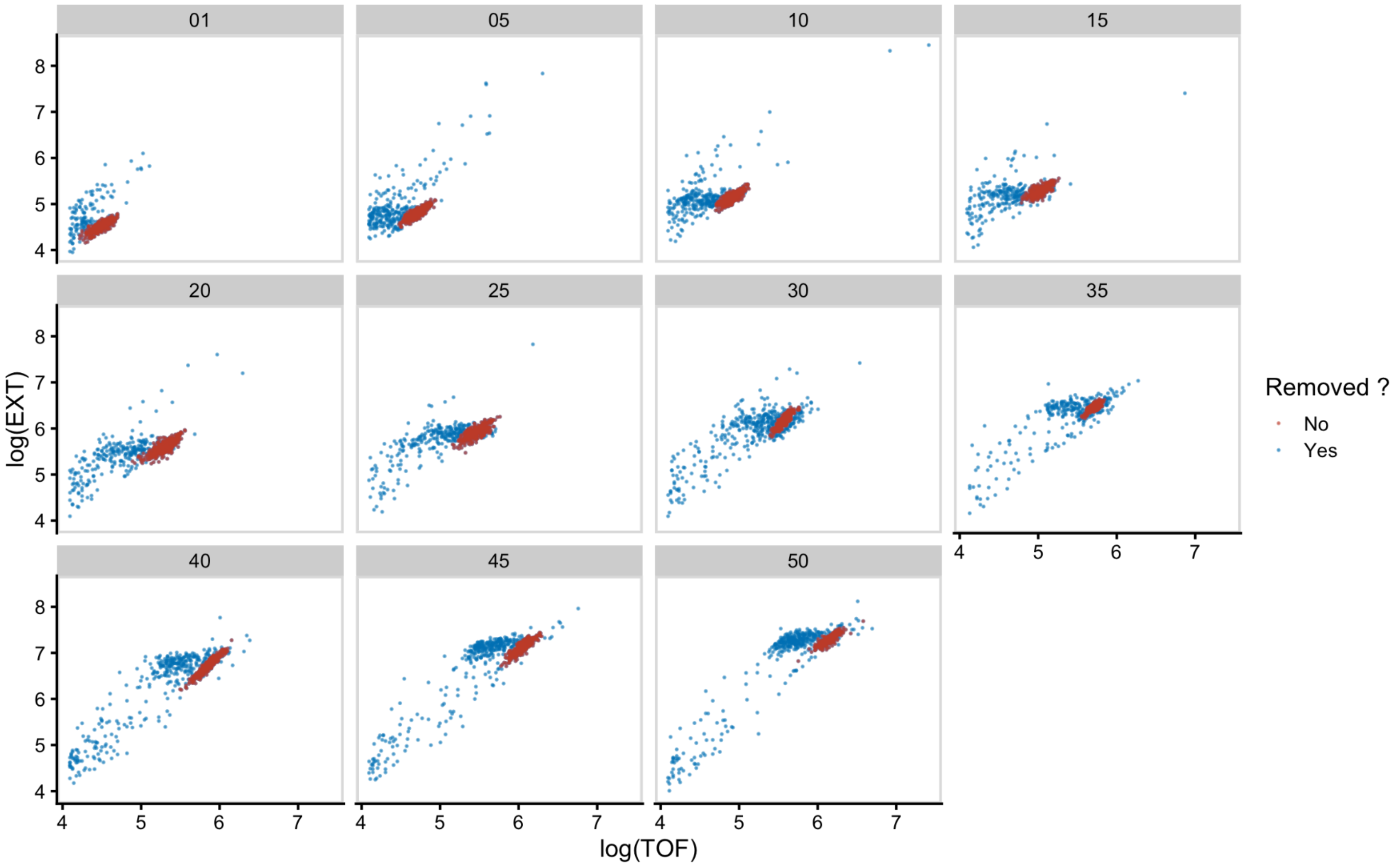


**S2 Fig. Mixture modeling of COPAS BIOSORT data was used to prune data.**
Mixture models of Gaussian distributions were fit to log transformed animal length (x-axis) and log transformed optical extinction (y-axis). Data from each hour of the experiment were separately analyzed and processed to remove clusters that did not include animal objects. Panels indicate experimental hours from which data were taken.

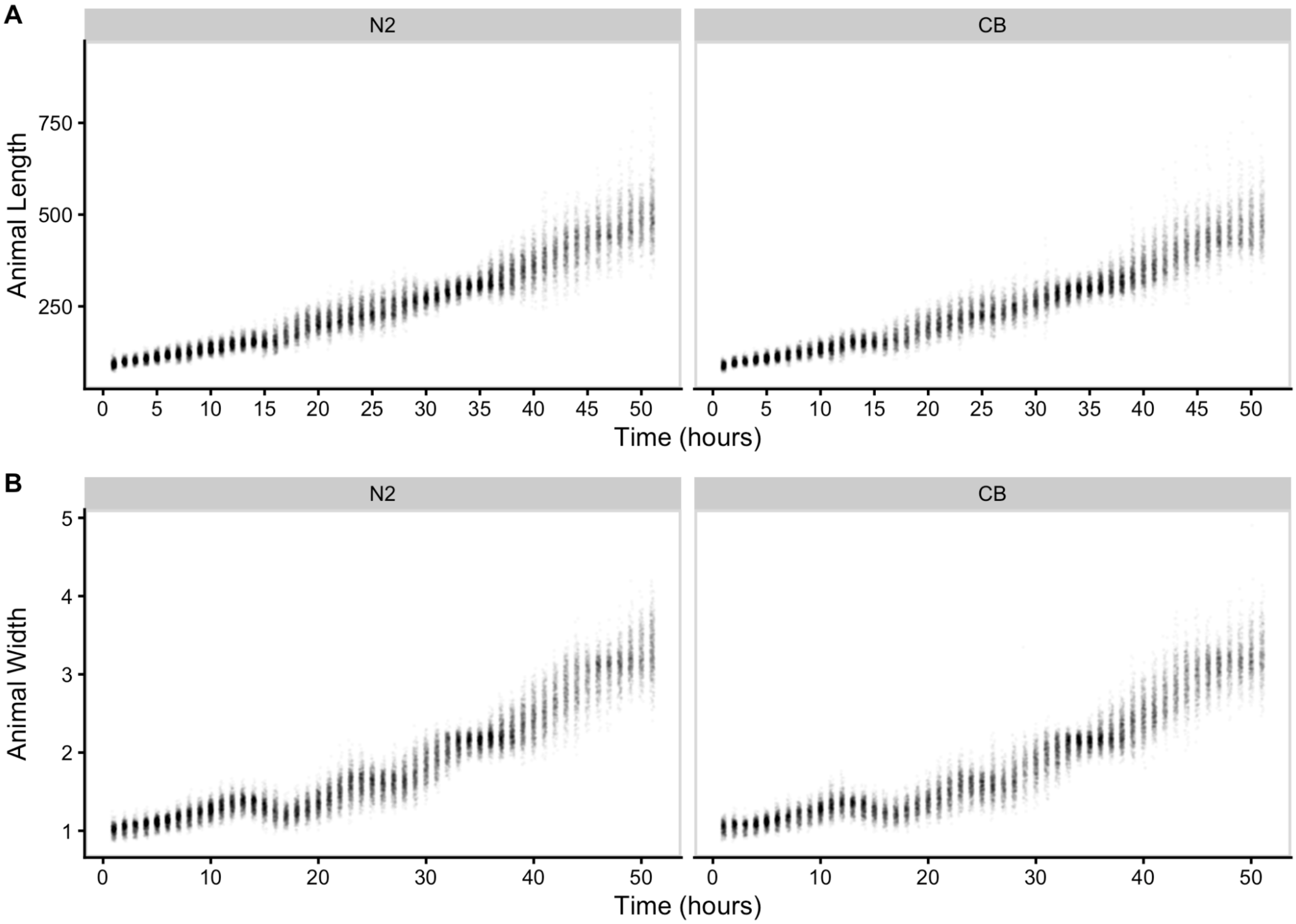


**S3 Fig. Pruned measurements of animal size.**
COPAS BIOSORT data of animal length (A) and width (B) after the removal of non-animal objects using model-based clustering methods. Each point represents an individual object that was measured.

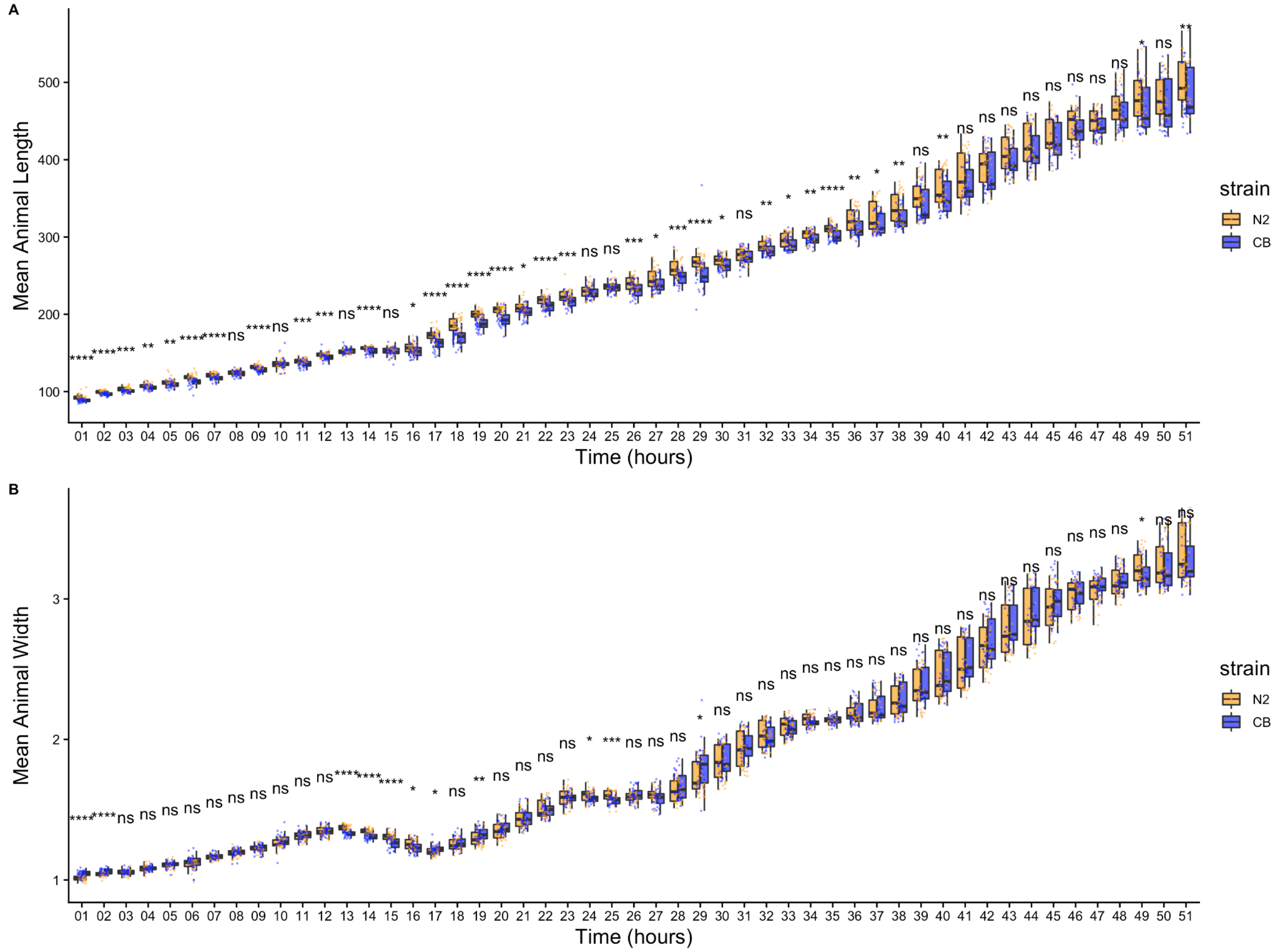


**S4 Fig. Comparison of means across developmental time points.**

Tukey boxplots of mean length **(A)** and mean width **(B)** for N2 (orange) and CB4856 (blue) over developmental time. The horizontal line in the middle of the box is the median, and the box denotes the 25th to 75th quantiles of the data. The vertical line represents the 1.5 interquartile range. Inset plots magnify mean animal size measurements from hour 48. Each point corresponds to the mean of a population of animals in each well. Statistical significance was calculated using a Wilcoxon test (ns = non-significant (p-value > 0.05); *, **, ***, and **** = significant (p-value < 0.05, 0.01, 0.001, or 0.0001, respectively).

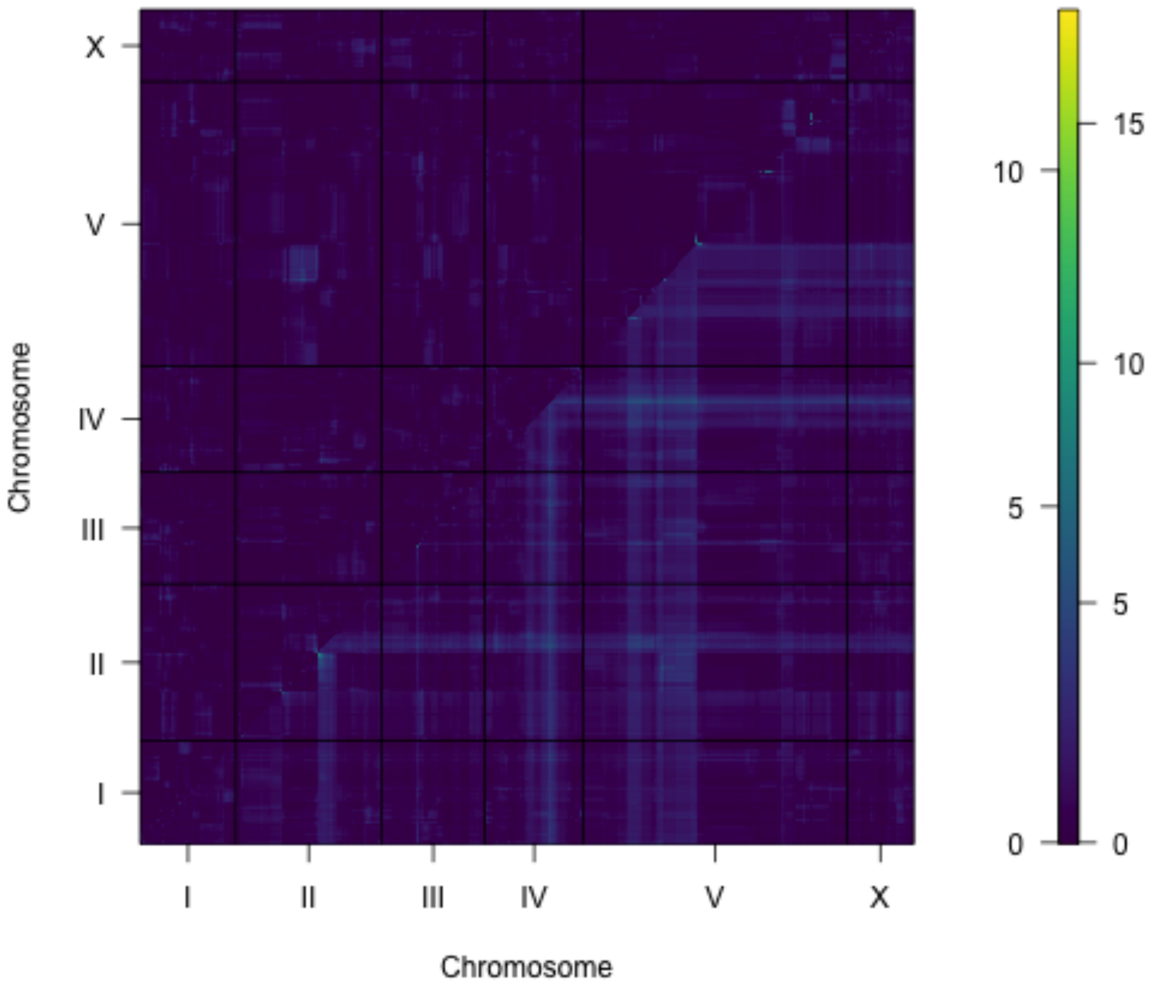


**S5 Fig. Two-dimensional genome scan for mean animal length.**
Log of the odds (LOD) scores are shown for each pairwise combination of loci, split by chromosome. The upper-left triangle contains the epistasis LOD scores and the lower-right triangle contains the LOD scores for the full model. LOD scores are colored, increasing from purple to green to yellow. The LOD scores for the epistasis model are shown on the left of the color scale and the LOD scores for the full model are shown on the right.

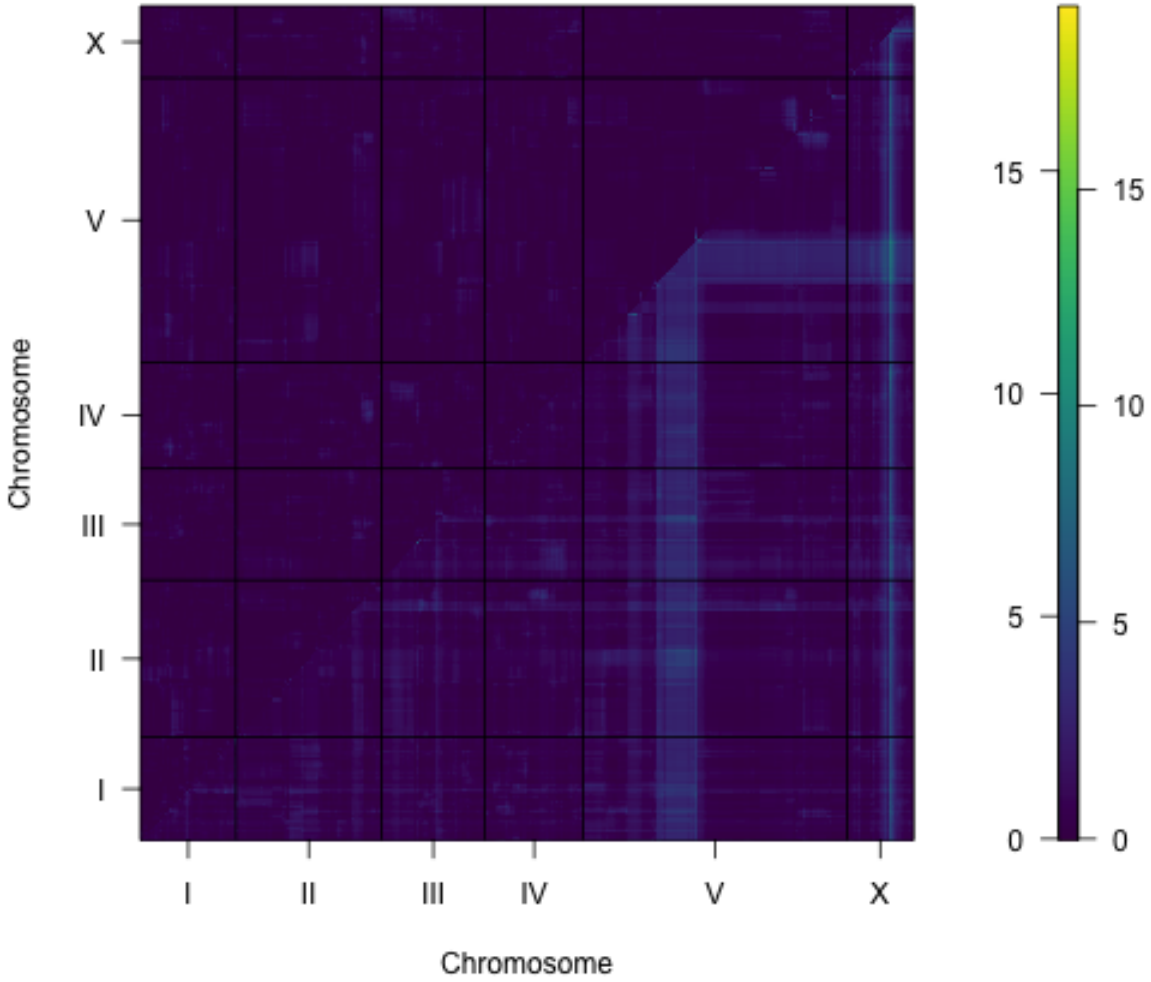


**S6 Fig. Two-dimensional genome scan for mean animal width.**
Log of the odds (LOD) scores are shown for each pairwise combination of loci, split by chromosome. The upper-left triangle contains the epistasis LOD scores and the lower-right triangle contains the LOD scores for the full model. LOD scores are colored, increasing from purple to green to yellow. The LOD scores for the epistasis model are shown on the left of the color scale and the LOD scores for the full model are shown on the right.

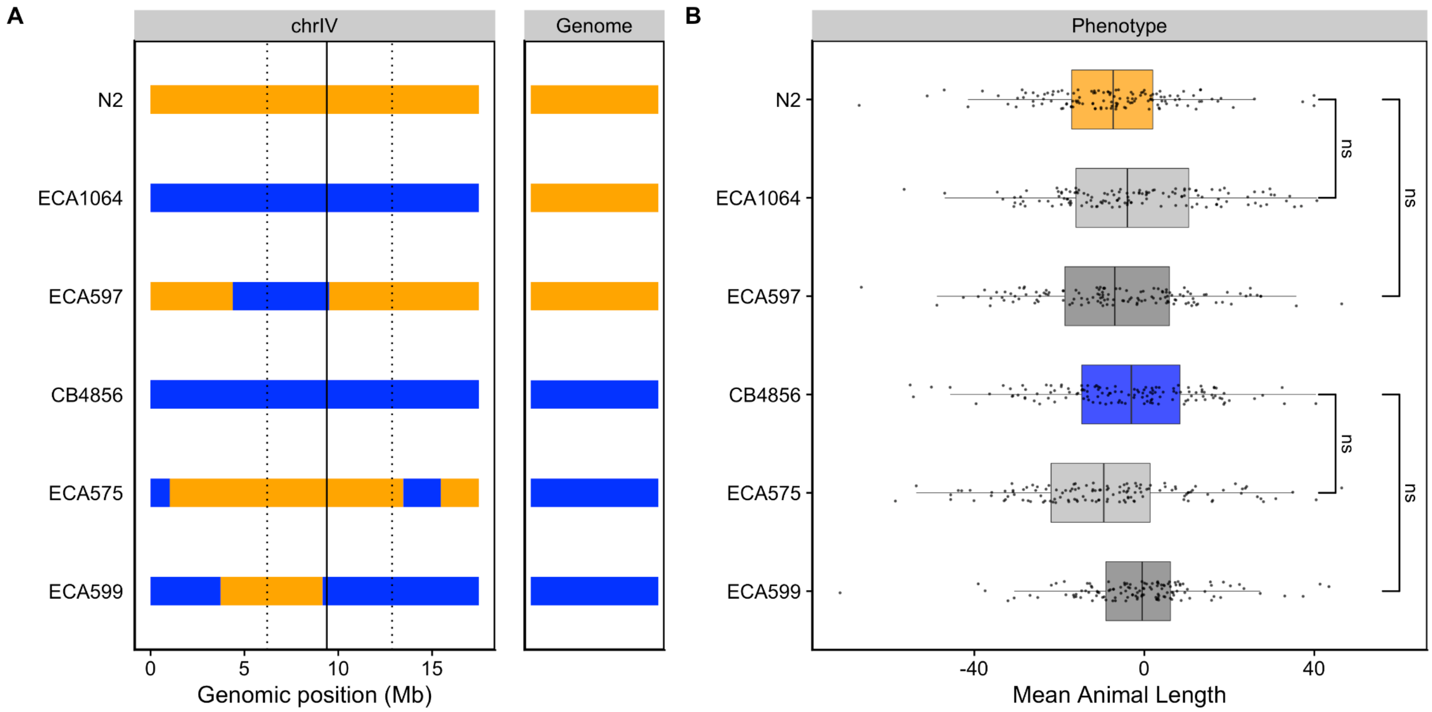


**S7 Fig. Validating the chromosome IV length-associated QTL.**

Strain genotypes are displayed as colored rectangles (N2: orange, CB4856: blue) for chromosome IV (left) and in general for the rest of the chromosomes (right). The solid vertical line represents the peak marker of the QTL. The dashed vertical lines represent the confidence interval. (B) Residual mean animal length (x-axis) is plotted as Tukey box plots against strain (y-axis). Each point corresponds to the mean length of a population of animals from each well in an assay. The boxes for the parental strains are colored: N2, orange; CB4856, blue. Statistical significance was calculated by Tukey's HSD (ns = non-significant, *p*-value > 0.05).

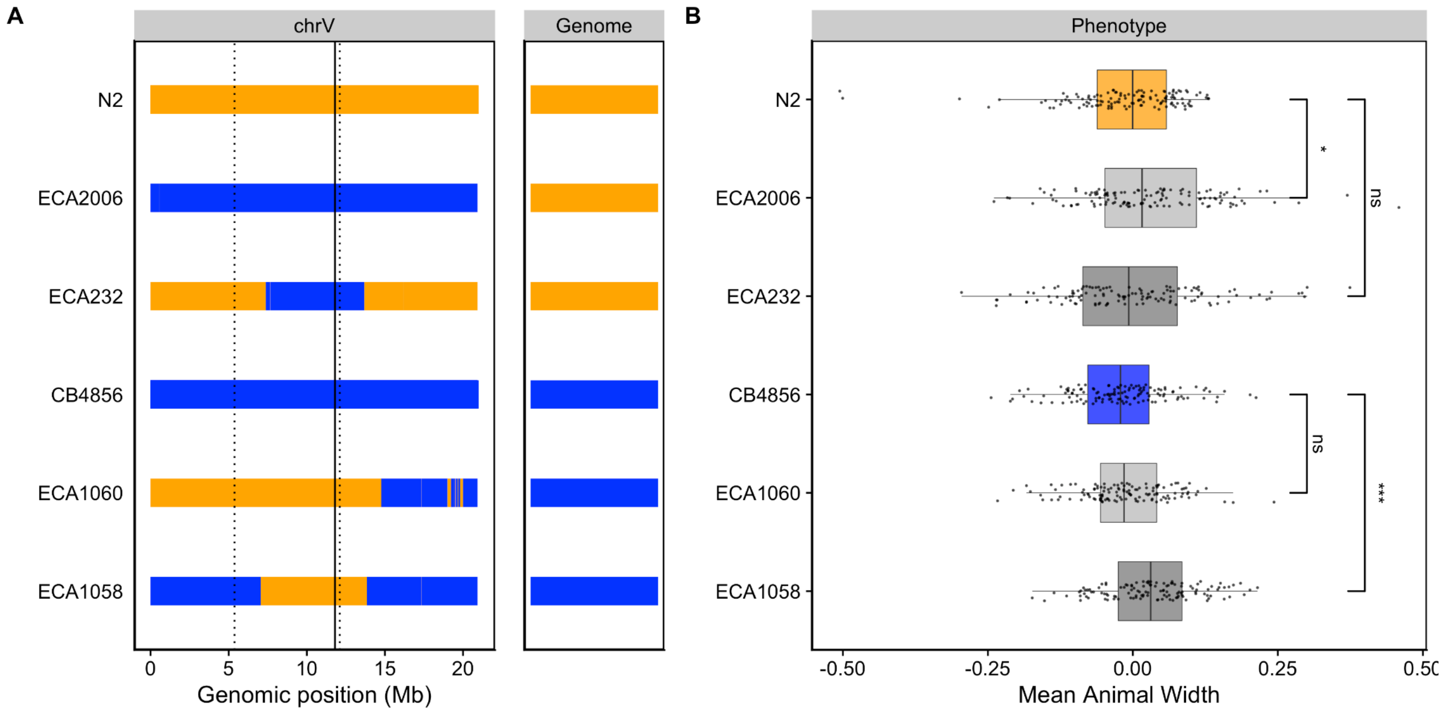


**S8 Fig. Validating the chromosome V width-associated QTL.**

Strain genotypes are displayed as colored rectangles (N2: orange, CB4856: blue) for chromosome V (left) and in general for the rest of the chromosomes (right). The solid vertical line represents the peak marker of the QTL. The dashed vertical lines represent the confidence interval. (B) Residual mean animal length (x-axis) is plotted as Tukey box plots against strain (y-axis). Each point corresponds to the mean width of a population of animals from each well in an assay. The parental strains are colored: N2, orange; CB4856, blue. Statistical significance was calculated by Tukey's HSD (ns = non-significant, *p*-value > 0.05; * and *** = significant, *p*-value < 0.05 or 0.001 respectively).

**S1 Text. Reagents used to generate CSSs and NILs.**

**List of RIAILs used:**

Set 2:

QX49, QX59, QX61, QX62, QX72, QX78, QX85, QX115, QX123, QX134, QX144, QX160, QX161, QX175, QX226, QX227, QX231, QX240, QX241, QX242, QX243, QX244, QX245, QX248, QX250, QX252, QX253, QX254, QX258, QX261, QX263, QX264, QX265, QX266, QX267, QX268, QX269, QX270, QX271, QX272, QX273, QX274, QX275, QX276, QX277, QX278, QX279, QX280, QX281, QX282, QX283, QX284, QX285, QX286, QX287, QX288, QX289, QX290, QX291, QX293, QX294, QX295, QX296, QX297, QX298, QX299, QX300, QX301, QX302, QX303, QX304, QX305, QX306, QX307, QX309, QX310, QX311, QX314, QX315, QX316, QX318, QX319, QX320, QX321, QX322, QX323, QX324, QX325, QX326, QX327, QX328, QX329, QX330, QX331, QX332, QX333, QX334, QX335, QX336, QX337, QX338, QX339, QX340, QX341, QX343, QX345, QX346, QX347, QX348, QX349, QX350, QX352, QX353, QX354, QX355, QX356, QX357, QX358, QX359, QX360, QX361, QX362, QX363, QX364, QX365, QX366, QX367, QX368, QX369, QX370, QX371, QX372, QX373, QX374, QX375, QX376, QX377, QX378, QX379, QX380, QX381, QX382, QX383, QX384, QX385, QX386, QX387, QX390, QX391, QX392, QX393, QX394, QX395, QX396, QX397, QX398, QX399, QX400, QX401, QX402, QX403, QX404, QX405, QX406, QX407, QX408, QX409, QX410, QX411, QX412, QX413, QX414, QX416, QX417, QX418, QX419, QX420, QX421, QX423, QX424, QX426, QX427, QX428, QX429, QX430, QX431, QX432, QX433, QX434, QX435, QX436, QX437, QX438, QX439, QX440, QX441, QX443, QX444, QX445, QX446, QX447, QX448, QX449, QX450, QX451, QX452, QX453, QX454, QX455, QX456, QX457, QX458, QX459, QX460, QX461, QX463, QX464, QX465, QX466, QX467, QX468, QX469, QX470, QX471, QX472, QX473, QX474, QX475, QX476, QX477, QX478, QX479, QX480, QX481, QX482, QX483, QX484, QX485, QX486, QX487, QX488, QX489, QX490, QX491, QX492, QX493, QX494, QX495, QX496, QX497, QX498, QX500, QX501, QX503, QX506, QX508, QX511, QX512, QX513, QX514, QX515, QX517, QX520, QX521, QX523, QX524, QX525, QX526, QX527, QX528, QX529, QX530, QX531, QX533, QX534, QX538, QX539, QX540, QX542, QX545, QX549, QX550, QX551, QX553, QX554, QX555, QX556, QX557, QX559, QX560, QX561, QX563, QX564, QX565, QX570, QX572, QX573, QX574, QX579, QX580, QX583, QX585, QX587, QX588, QX594, QX596, QX597, QX598

**Reagents to generate CSSs and NILs:**

| Strain | Genotype | Constructed from | Left primer | Right primer |
|---|---|---|---|---|
| ECA232 | *eanIR152*[V, CB4856>N2] | QX450xN2 | oECA799 & oECA800 | oECA745 & oECA746 |
| ECA575 | *eanIR324*[IV, N2>CB4856] | N2xCB4856 | oECA1132 & oECA1133 | oECA1135 & oECA1136 |
| ECA597 | *eanIR330*[IV, CB4856>N2] | ECA231xN2 | oECA781 & oECA782 | oECA857 & oECA858 |
| ECA599 | *eanIR332*[IV, N2>CB4856] | ECA598xCB4856 | oECA781 & oECA782 | oECA857 & oECA858 |
| ECA828 | *eanIR359*[X, N2>CB4856] | N2xCB4856 | oECA1313 & oECA1314 | oECA1246 & oECA1247 |
| ECA929 | *eanIR411*[X, CB4856>N2] | N2xCB4856 | oECA1313 & oECA1314 | oECA1246 & oECA1247 |
| ECA1058 | *eanIR433*[V, N2 > CB4856] | ECA1029xCB4856 | oECA1408 & oECA1409 | oECA1341 & oECA1342 |
| ECA1060 | *eanIR435*[V, N2 > CB4856] | ECA554xCB4856 | oECA745 & oECA746 | oECA763 & oECA764 |
| ECA1064 | *eanIR439*[IV, CB4856>N2] | N2xCB4856 | oECA1131 & oECA1132 | oECA1135 & oECA1136 |
| ECA2006 | *eanIR446*[V, CB4856>N2] | N2xCB4856 | oECA1141 & oECA1142 | oECA1147 & oECA1148 |

**Primers:**

| Primer | Genomic position | Sequence |
|---|---|---|
| oECA745 | V:13,110,045 | tgcagaggtggagtaaccct |
| oECA746 | V:13,110,045 | ctcggtctctcccccactaa |
| oECA763 | V:15,121,356 | cgcacattctttatttctggcg |
| oECA764 | V:15,121,356 | atcggccgtttttcacctga |
| oECA781 | IV:5,110,734 | gagcactttggcgactttcg |
| oECA782 | IV:5,110,734 | tccgggcaaattagtgtggc |
| oECA799 | V:7,862,556 | ttctcgctactggaacacgc |
| oECA800 | V:7,862,556 | tcaagaagcgttgggaagtct |
| oECA1131 | IV:1,039,851 | tacccaccgcatcaaaacca |
| oECA1132 | IV:1,039,851 | acaggcgttcaaagacacca |
| oECA1135 | IV:17,317,014 | tttcagacaggaaagcgcct |
| oECA1136 | IV:17,317,014 | gttgagagatccggaccgac |
| oECA1141 | V:144,547 | ctcatgggagtaacctgggc |
| oECA1142 | V:144,547 | cggtgacaacggagaatcca |
| oECA1147 | V:20,622,851 | gtttagtaccagcggggcat |
| oECA1148 | V:20,622,851 | tgcattccgacccaagagac |
| oECA1246 | X:11,696,902 | tgcggtgggacttttcttgt |
| oECA1247 | X:11,696,902 | gtcccagcatgtaaccgtct |
| oECA1313 | X:8,038,337 | gctgtgcaggactggatgta |
| oECA1314 | X:8,038,337 | tgctttctgatctgtgccgt |
| oECA1341 | V:7,104,674 | cccatccccacaatgtttcg |
| oECA1342 | V:7,104,674 | aatcgacgagtggcacttgt |
| oECA1408 | V:3,778,859 | cacgtgccctttttgcaatga |
| oECA1409 | V:3,778,859 | gagctcccggaaaactcgaa |
